# A robust ultrasensitive transcriptional switch in noisy cellular environments

**DOI:** 10.1101/2023.08.15.553401

**Authors:** Eui Min Jeong, Jae Kyoung Kim

## Abstract

Ultrasensitive transcriptional switches enable sharp transitions between on and off states of transcription and are essential for cells to respond to environmental cues precisely. However, conventional switches, relying on direct repressor-DNA binding, are extremely sensitive to noise. Here, we discovered an alternative design combining three indirect transcriptional repression mechanisms, sequestration, blocking, and displacement, to generate a noise-resilient ultrasensitive switch. Although sequestration alone can generate an ultrasensitive switch, it remains sensitive to noise because the unintended transcriptional state induced by noise can persist for long periods. However, by jointly utilizing blocking and displacement, these noise-induced transitions can be rapidly restored to the original transcriptional state. Because this transcriptional switch is effective in noisy cellular contexts, it goes beyond previous synthetic transcriptional switches utilizing direct repression mechanisms, making it particularly valuable for robust synthetic system design. Our findings also provide insights into the evolution of robust ultrasensitive switches in real cells. Specifically, the concurrent use of seemingly redundant indirect repression mechanisms in diverse biological systems appears to be a strategy to achieve noise-resilience of ultrasensitive switches.

**GRAPHICAL ABSTRACT:** 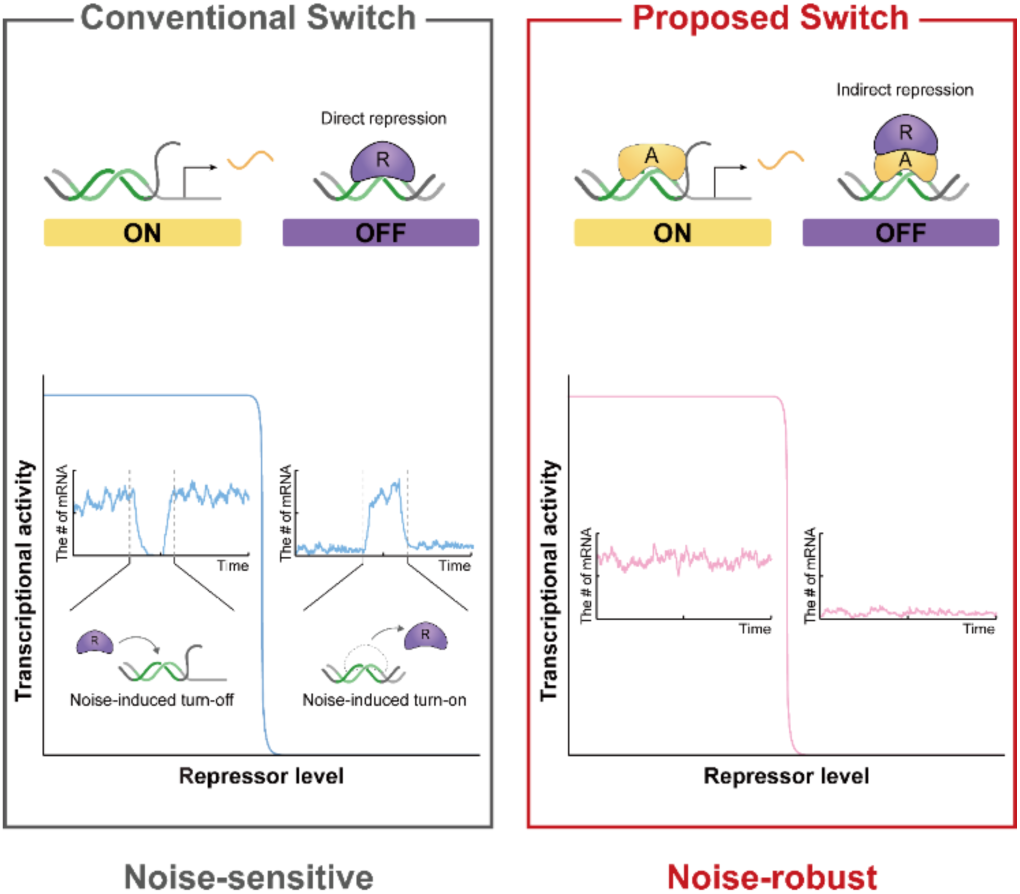

## INTRODUCTION

Ultrasensitive responses in biological systems are input/output relationships that exhibit a highly sensitive response to changes in input or stimulus (1). For example, transcriptional ultrasensitive switches are turned on or off depending on whether the level of the transcription factor exceeds or falls below a threshold. This distinctive property renders these switches valuable for systems requiring precise decision-making contingent on the number of input transcription factors. Moreover, these switches serve as catalysts for diverse cellular functions including signal amplification and generation of bistability and oscillations (2,3), thereby bestowing them with versatile utility across various biological contexts. An ultrasensitive transcriptional switch can be generated by the cooperative binding of transcription factors to multiple DNA binding sites (2-4) or their titration via decoy sites (5). And these mechanisms are used in the majority of synthetic transcriptional switches (6,7). However, recent studies have raised concerns about the effectiveness of these conventional transcriptional switches in noisy cellular environments (8-10). Specifically, when noise induces an undesired transition between the transcriptional ‘on’ and ‘off’ irrespective of the concentration of the transcription factor, this can persist for long periods.

Alternatively, transcriptional repressors can indirectly suppress transcription by targeting transcriptional activators rather than directly binding to DNA (11,12). This suppression can occur by three different mechanisms. One mechanism is when repressors sequester transcriptional activators, preventing their binding to DNA. For example, sigma factors or basic leucine zippers can be sequestered through interactions with anti-sigma factors or inhibitors, respectively (13-15). A second mechanism is when repressors bind to activators already bound to DNA, leading to transcriptional inhibition. This blocking is utilized when PHO80, a kinase component in a signaling pathway of yeast, inhibits PHO4 (16). In addition to sequestration and blocking, the third mechanism is when repressors displace DNA-bound activators. An illustrative case is the IκB protein, which reduces NF-κB activator’s binding affinity with DNA, displacing it from the DNA (17). We recently found that the combination of these three mechanisms, sequestration, blocking and displacement, can generate ultrasensitive transcriptional switches (18). Interestingly, these three indirect repression mechanisms synergistically function in systems like the NF-κB oscillator and circadian clocks (17,19-23), where precise ultrasensitivity against noise is crucial. This led us to hypothesize that an ultrasensitive switch based on the combination of multiple indirect repression mechanisms could be robust against noise unlike the conventional switches relying on direct repression.

Here, we tested that hypothesis and indeed found that the ultrasensitive switch based on the combination of the sequestration, blocking, and displacement can stably sustain the transcriptional ‘on’ and ‘off’ states even in the presence of noise. Specifically, our research unveils that the combined integration of these three repression mechanisms immediately restores the original ‘off’ and ‘on’ states even when noise triggers transition to the undesired transcriptional state. In this switch, the sequestration generates a sharp transition from the transcriptional ‘on’ state to the ‘off’ state as the molar ratio between repressors and activators shifts from below one to above one. Furthermore, even if the transcriptional ‘off’ state transits to the ‘on’ state due to noise, the blocking rapidly restores the original ‘off’ state when the molar ratio surpasses one. In addition, the displacement ensures that the noise-induced transcriptional ‘on’ state swiftly returns to the original ‘off’ state when the molar ratio falls below one. This finding is supported by further analysis of identified mutation disrupting the displacement in the mammalian circadian clock. Taken together, we propose an ultrasensitive transcriptional switch that operates effectively even in noisy cellular environments and offers a promising strategy for designing robust synthetic switches.

## MATERIAL AND METHODS

### Calculations of the transcriptional activity and the Fano factor

We calculated the transcriptional activity and the Fano factor of mathematical models with an approach established by A. Sanchez et al (8). The equations resulting from this analysis are presented in Table S1, while the detailed derivation process is provided in the Supplementary Information.

## RESULTS

### Noise triggers undesirable activation and repression of the cooperative binding-based switch

Ultrasensitivity can be generated through the cooperative binding of transcriptional repressors to multiple DNA sites (3,24). To investigate this mechanism, we used a previously developed model of a cooperative binding-based switch (8) (Figure 1A). In this model, the DNA has three sites that can bind to the repressor (*R*) with a dissociation constant of K_r_. When one site of the DNA is occupied, *R* can bind to the other two sites with a dissociation constant of *cK*_*r*_. Similarly, when two sites of the DNA are occupied, then *R* can bind to the remaining site with a dissociation constant of *c*^2^*K*_*r*_. Therefore, when *c* = 1, *R* binds independently to each of the three sites, but when *c* < 1, *R* binds more favorably to the DNA when one or two sites are already occupied than when all sites are unoccupied (i.e., *R* cooperatively binds). Given that when all three sites are occupied by *R*, transcription is suppressed (Figure 1A, gray box), the transcriptional activity is the probability that at least one site is unoccupied by *R* at the steady state. This probability can be calculated from the chemical master equation (CME) governing the system with respect to the effective total number of the repressor 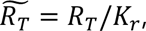 where *R*_*T*_ is the total number of repressors (Table S1; see the Materials and Methods).

**Figure 1.**
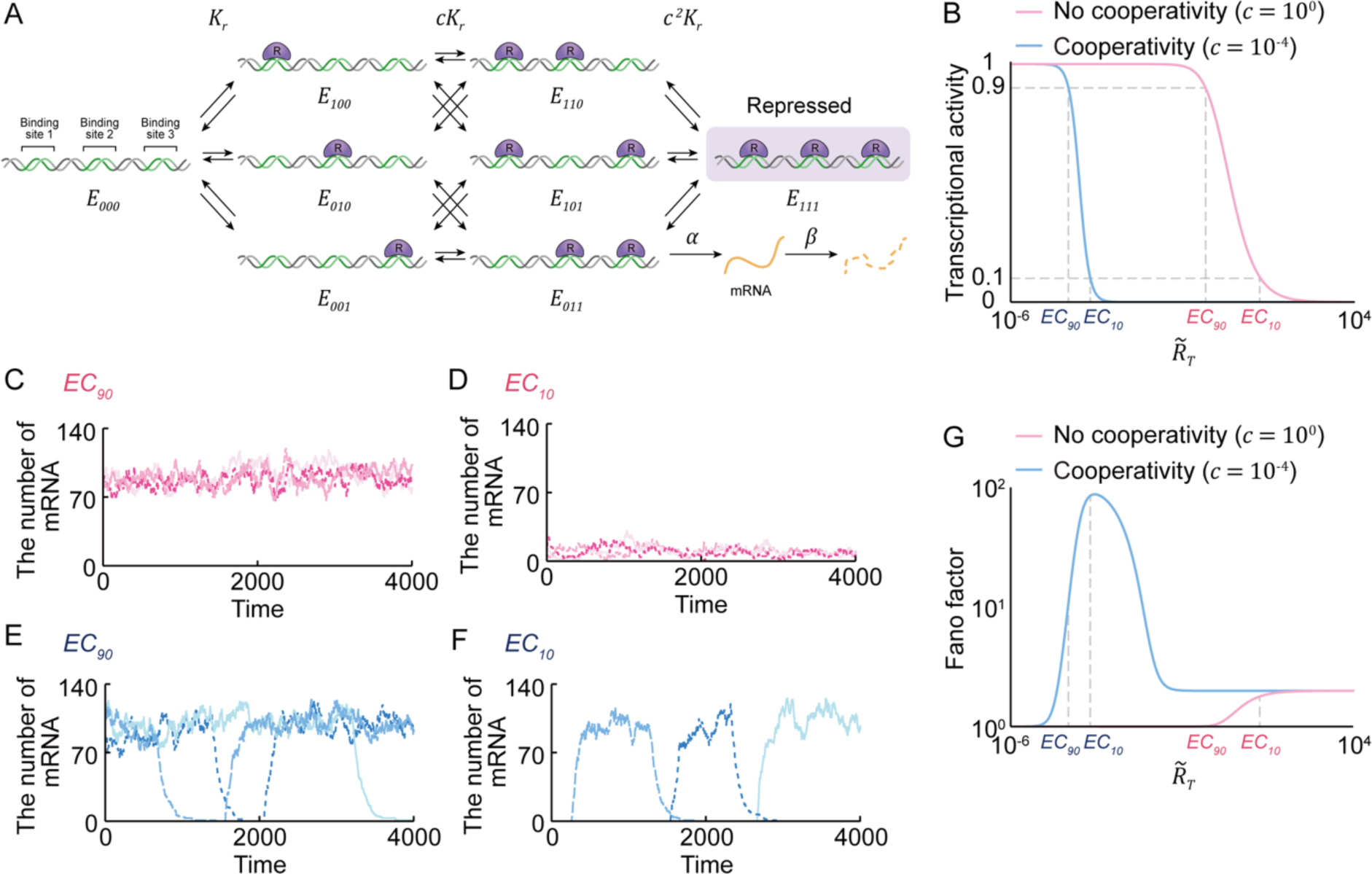
The transcriptional ultrasensitive switch based on cooperative binding suffers from undesired switch transition due to noise. (**A**) Schematic diagram of the model describing the binding of the repressor (*R*) to three sites on DNA. *R* can bind to an unoccupied site (*E*_000_) with a dissociation constant of *K*_*r*_. When one site is occupied (*E*_001_, *E*_010_, or *E*_100_), *R* can bind to the other two sites with a dissociation constant of *cK*_*r*_. When two sites are occupied (*E*_110_, *E*_101_, or *E*_011_), *R* can bind to the remaining site with a dissociation constant of *c*^2^*K*_*r*_. Accordingly, when *c* < 1, cooperativity is present, while absent when *c* = 1. When all sites are occupied (*E*_111_), the transcription is repressed (gray box); otherwise, mRNA is transcribed with the rate of *α* and decays with the rate of *β*. (**B**) Consequently, as the total effective number of repressors 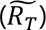 increases, the probability that transcription is activated (i.e., transcriptional activity) decreases, and the system transitions from a transcriptional activation phase (TAP) to a transcriptional repression phase (TRP). This transition in the presence of cooperativity (blue line) is more sensitive to changes in 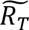 (i.e., a shorter distance between *EC*_90_ and *EC*_10_) compared to in the absence of cooperativity (red line). (**C – D**) In the absence of cooperativity, the simulated timeseries of mRNA fluctuate around their high and low averages during TAP (i.e., when 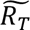 is at *EC*_90_) (C) and during TRP (i.e., when 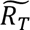 is at *EC*_10_) (D), respectively. (**E – F**) On the other hand, in the presence of cooperativity, the number of mRNAs can decrease to 0 even during TAP and can significantly increase even during TRP (F). (**G**) As a result, in the presence of cooperativity, the number of mRNAs exhibits greater variance, resulting in a higher Fano factor compared to in the absence of cooperativity.

The transcription is turned on with a high probability when 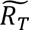 is low, referred to as the transcriptional activation phase (TAP). As 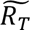 increases, the probability decreases to zero, reaching the transcriptional repression phase (TRP; Figure 1B). The sharpness of this transition from transcriptional activation during TAP to repression during TRP depends on the cooperativity *c*. Specifically, when there is no cooperativity (i.e., *c* = 1), the transition shows a similar sensitivity with the Michaelis-Menten equation (Figure 1B, red line). In contrast, when repressors bind cooperatively (i.e., *c* < 1), the sensitivity of the transition increases (Figure 1B, blue line). This sensitivity can be quantified using the effective Hill coefficient log 81 / log(*EC*_10_/*EC*_90_), where *EC*_90_ and *EC*_10_ are the values of 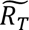 at which the transcriptional activity becomes 0.9 and 0.1, respectively (1). The measured effective Hill coefficients in the presence and absence of cooperativity are about 3 and 1.3, respectively (Figure 1B).

Next, we simulated the time series of mRNA (how the number of mRNAs produced by transcription fluctuates over time) during TAP (transcriptional activation) and TRP (repression) using the Gillespie algorithm. We chose the values of 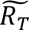 (total number of repressors) at *EC*_90_ for TAP and at *EC*_10_, for TRP. In the absence of cooperative binding between repressors, mRNA numbers slightly fluctuate around their high average during TAP (Figure 1C) and around their low average during TRP (Figure 1D). However, in the presence of cooperativity, mRNA numbers often drop to zero even during TAP (Figure 1E), and conversely often lift up to high levels even during TRP (Figure 1F). These undesired transcriptional states, arising due to noise, persist because transition between the transcriptional activation and repression occurs rarely due to cooperative binding (8). As a result, cooperativity increases the variance in the number of mRNAs transcribed, yielding a much higher Fano factor compared to the absence of cooperativity (Figure 1G and Table S1). This shows that cooperative binding-based transcriptional switches, which induce transcriptional repression through direct binding to DNA, can cause undesirable transitions of transcriptional states that persist for a long time.

### Noise turns on the sequestration-based switch during TRP

Transcriptional repression can occur not only through direct binding to DNA, but also by indirectly inhibiting the transcriptional activator that promotes transcription. Among the indirect repression mechanisms, sequestration of transcriptional activators preventing their binding to DNA has been identified to generate ultrasensitivity (25-28). To investigate whether this type of switch is robust to noise, we constructed a model describing the sequestration (Figure 2A). In this model, the free activator (*A*) binds to the free DNA (*E*_*A*_) with a dissociation constant of *K*_*a*_ to form the activated DNA (*E*_*A*_). Transcription is inhibited when the free repressor (*R*) binds to *A* to form an inactive complex (*R*_*A*_) with a dissociation constant of *K*_*s*_ (i.e., sequestration; Figure 2A, gray box). Thus, the transcriptional activity is defined as the probability that the DNA is activated by the binding of *A*, and not sequestered by *R* at the steady state. We derived the probability as a function of 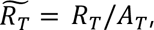 the molar ratio between total numbers of the activator (*A*_*T*_ = *A* + *R*_*A*_ + *E*_*A*_) and the repressor (*R*_*T*_ = *R* + *R*_*A*_) (Table S1; see Materials and Methods for details).

**Figure 2.**
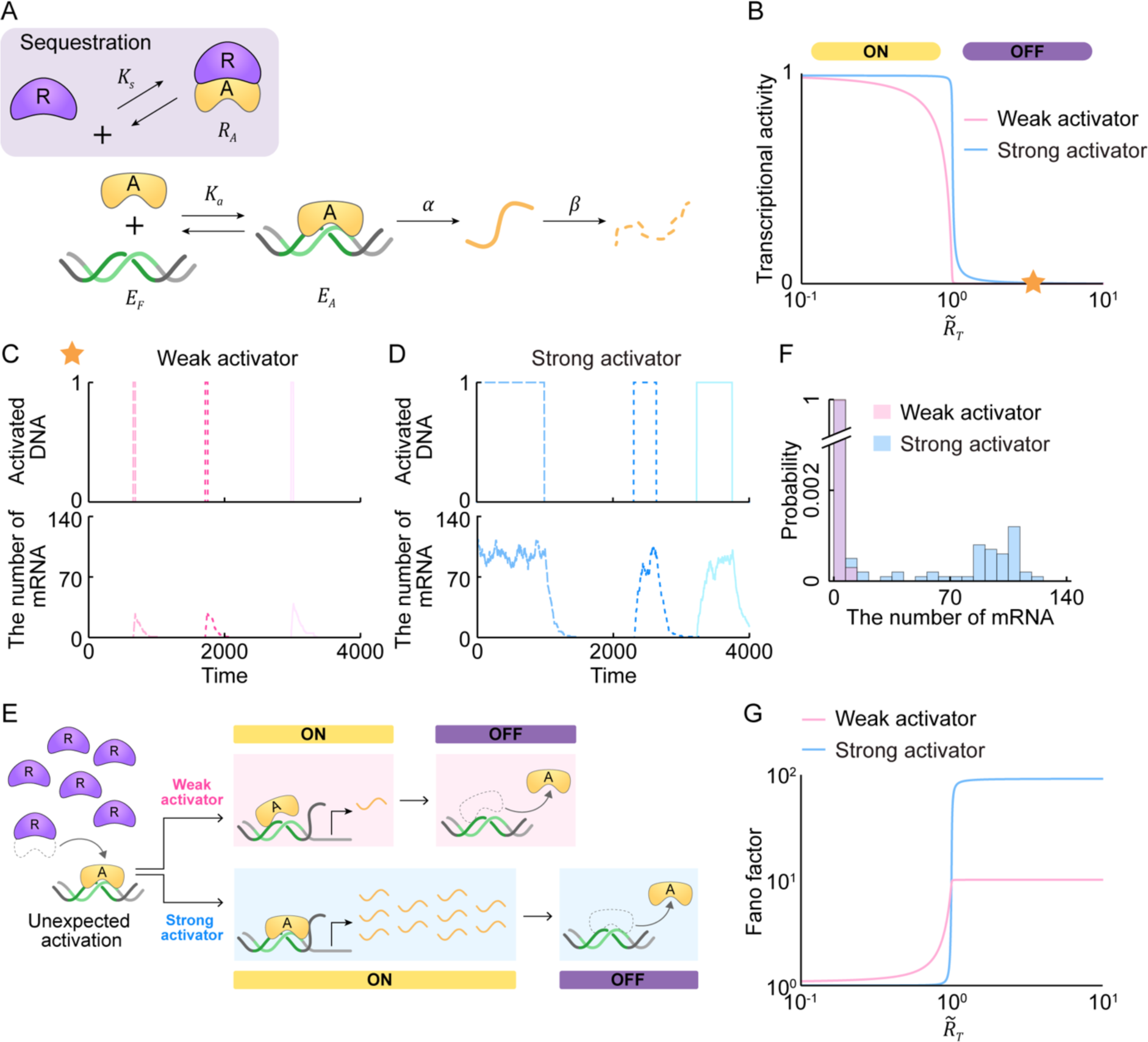
A transcriptional ultrasensitive switch based on solely on sequestration is sensitive to noise during TRP due to undesired transcriptional activation. (**A**) Schematic diagram of the model describing the sequestration. Binding of the activator (*A*) to the DNA (*E*_*A*_) with a dissociation constant of *K*_*a*_ forms the activated DNA (*E*_*A*_) to promote the transcription of mRNA with the rate of *α*, which decays with the rate of *β*. On the other hand, binding of the repressor (*R*) to *A* with a dissociation constant of *K*_*s*_ leads to transcriptional repression (i.e., sequestration; gray box). (**B**) Consequently, when the molar ratio 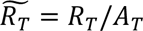 between the total activator (*A*_*T*_) and repressor (*R*_*T*_) is less than one (TAP), unsequestered activators can turn on transcription. In contrast, when the molar ratio exceeds one (TRP), almost all activators are sequestered by repressors, and thus transcription is turned off. As a result, around 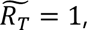 a sensitive transition from TAP to TRP occurs with respect to changes in 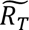. Notably, strong activators, which bind tightly to DNA, exhibit a more sensitive transition compared to weak activators that bind loosely. (**C – D**) The undesired activation is restored during TRP to the repressed state for a weak activator (C), unlike a strong activator (D). Here, the timeseries of activated DNA and mRNA are simulated at 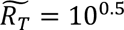 (orange star in (B)). (**E**) When noise-induced release of *A* from the *R*_*A*_ complex occurs, transcription can be turned on by binding of the *A* to DNA (i.e, unexpected activation occurs). Such undesired activation during TRP persists for a long time with a strong activator due to tight binding (lower arrow), but not with a weak activator (upper arrow). This sharply increases the number of mRNAs. (**F**) As a result, the stationary distribution of mRNA molecules shows an additional peak far from zero, leading to bimodality with a strong activator (blue bars) unlike with a weak activator (red bars). (**G**) Such bimodality deviates the distribution from a Poisson distribution, resulting in a higher Fano factor for a strong activator compared to a weak activator during TRP.

The transcription turns on and off, respectively, when the molar ratio 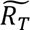 is either below (TAP) or above one (TRP). The transition between transcriptional activation during TAP and repression during TRP becomes more sensitive as the binding between *A* and the DNA becomes stronger (i.e., 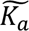 decreases; Figure 2B). This is because a strong activator (i.e., small 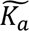) can promote transcription even when it is present in small numbers, sustaining transcriptional activation during TAP as long as there are more activators than repressors (i.e., 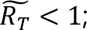 Figure 2B, blue line). However, when the molar ratio exceeds one (i.e., 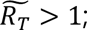 Figure 2B), the majority of *A* is sequestered by *R*. As a result, a sensitive transition from TAP to TRP occurs near 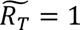 with the high effective Hill coefficient of about 50. On the other hand, the transition occurs gradually with the effective Hill coefficient of about 2 in the case of a weak activator (i.e., large 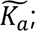 Figure 2B, red line).

Next, we simulated the timeseries of activated DNA and mRNA at the value of 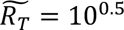 in TRP (Figure 2B, orange star). For a weak activator, the activated DNA indued by noise is swiftly restored to the repressed state, keeping the number of mRNAs consistently low around zero (Figure 2C). On the other hand, for a strong activator, this undesired activation of DNA during TRP persists over a long period, leading to a significant increase in the number of mRNAs (Figure 2D). This disparity arises when a sequestered *A* is released from *R* and binds to the DNA despite the system being in the TRP (leading to undesired activation; Figure 2E). Subsequently, a weak activator would quickly dissociate from the DNA and thus transcription is turned off to its desired state (Figure 2E, upper arrow). Meanwhile, a strong activator would remain bound to the DNA for a long time (Figure 2E, lower arrow). As a result, the undesired transcriptional activation persists even during TRP, resulting in a considerable increase in the number of mRNAs. This leads to a bimodal distribution of mRNA molecules deviating substantially from a Poisson distribution, with a strong activator (Figure 2F, blue bars) unlike with a weak activator (Figure 2F, red bars). Thus, a strong activator yields much higher transcriptional noise compared to a weak activator during TRP (Figure 2G). In conclusion, when the binding between the activator and DNA is tight, a sensitive transcriptional response can be generated with sole sequestration. However, this tight binding leads to high transcriptional noise because undesired transcriptional activation during TRP persists for a long period.

### Noise turns off a sequestration- and blocking-based switch during TAP

When transcription is regulated by sequestration alone, high transcriptional noise occurs during TRP due to the absence of repression for the activator bound to the DNA. This DNA-bound activator can be suppressed when the repressor blocks transcription either by forming a complex with the DNA-bound-activator (i.e., blocking), or by pulling off the activator from the DNA (i.e., displacement). Indeed, these repression mechanisms are commonly employed in addition to the sequestration mechanism in many biological systems (11,18). We first added blocking to the sole sequestration model, where the repressor (*R*) can directly bind to the DNA-bound-activator (*E*_*A*_) with a dissociation constant of *K*_*b*_ to form the repressed DNA (*E*_*R*_; Figure 3A, gray box). Based on this model, we derived the transcriptional activity as a function of the molar ratio 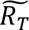 (Table S1; see Materials and Methods for details). The derived transcriptional activity exhibits a high sensitivity of response to changes in 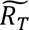 (i.e., the effective Hill coefficient of about 50; Figure 3B). Furthermore, the addition of blocking rapidly restores the undesired activation of DNA during TRP to the repressed state (Figure 3C, top panel). This is evident in the simulated timeseries of mRNA, which remains close to zero (Figure 3C, bottom right panel) without the sharp increase of mRNAs seen with sequestration alone (Figure 3C, bottom left panel). That is, the undesired transcription triggered by the noise-induced activator binding to DNA during TRP is immediately inhibited by the repressor via blocking (Figure 3D, lower arrow) in contrast to sole sequestration (Figure 3D, upper arrow). As a result, in the presence of blocking, the distribution of mRNA molecules is concentrated near zero (Figure 3E, blue bars) rather than bimodal (Figure 3E, red bars), reducing the transcriptional noise during TRP (Figure 3F and Table S1).

**Figure 3.**
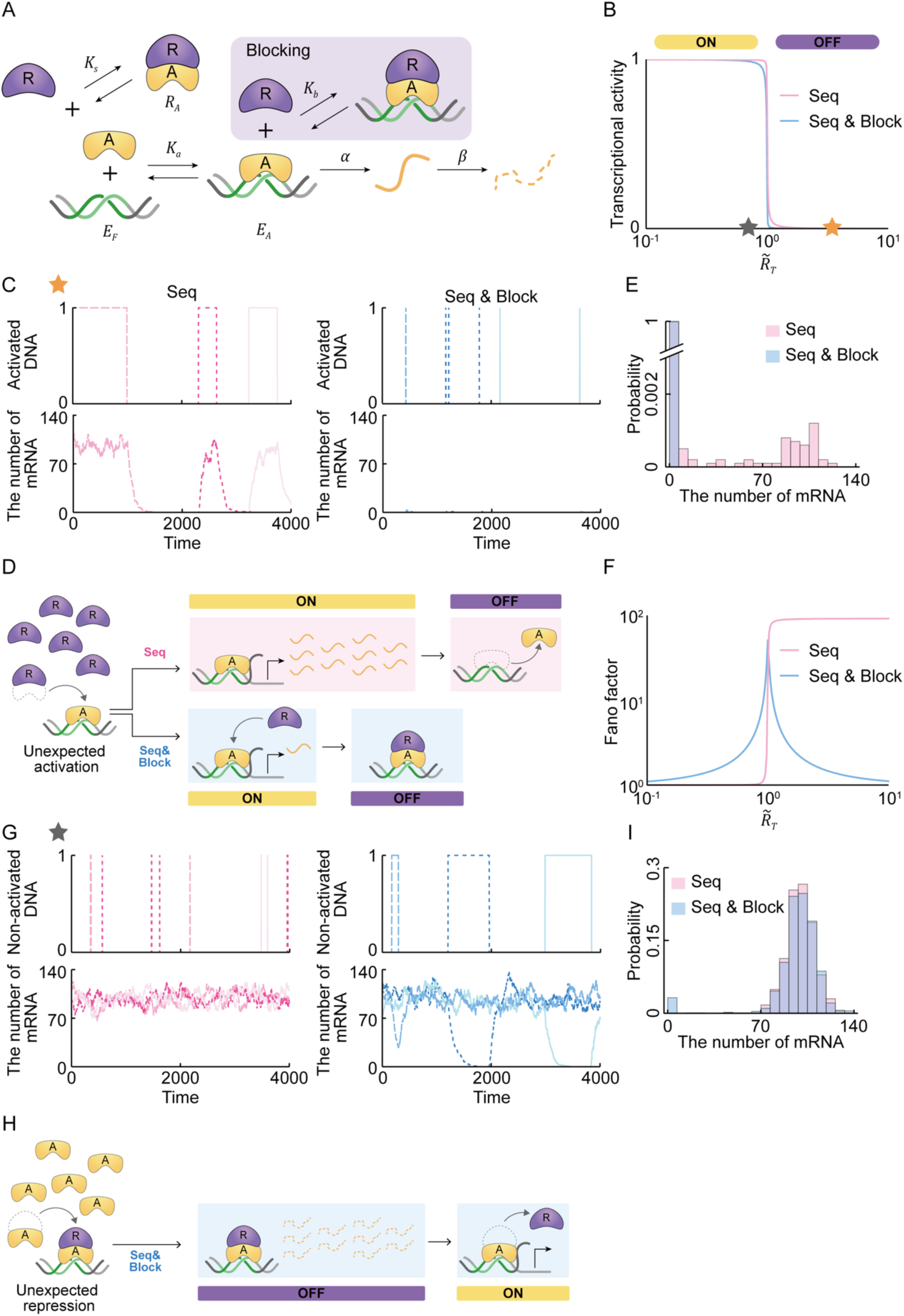
A transcriptional ultrasensitive switch based on the combination of sequestration and blocking is sensitive to noise during TAP due to undesired transcriptional repression. (**A**) A blocking mechanism is added to the sole sequestration model. The repressor (*R*) binds to the DNA-bound-activator (*E*_*R*_) with a dissociation constant of *K*_*d*_ to block transcription (i.e., blocking; gray box). (**B**) The combination of sequestration and blocking (blue line) generates an ultrasensitive switch similar to sequestration alone (red line). (**C**) However, unlike sole sequestration, the combination of sequestration and blocking rapidly restores the undesired activation of DNA during TRP to the repressed state. (**D**) When unwanted binding of activator to DNA occurs during TRP, the repressor rapidly inhibits the activator in the presence of blocking (lower arrow), but not in its absence (upper arrow). (**E**) As a result, the stationary distribution of mRNA molecules is concentrated around zero (blue bars) without bimodality observed in sole sequestration (red bars). (**F**) Thus, the noise level dramatically reduces during TRP in the presence of blocking. However, the addition of blocking to sequestration increases the noise level during TAP. (**G**) Such high transcriptional noise is due to the persistence of undesired repression for long periods even during TAP when blocking is added. Here, the timeseries of non-activated DNA (i.e., free or repressed DNA) and mRNA are simulated at 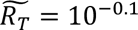 (gray star in (B)). (**H**) When noise-induced binding of *R* to DNA-bound-activator occurs (i.e., blocking occurs), due to their tight binding, the undesired repression during TAP persists for a long time. This leads to the sharp decrease of mRNAs close to zero. (**I**) As a result, the stationary distribution of mRNA molecules shows an additional peak around zero, leading to bimodality with blocking (blue bars) unlike with sequestration alone (red bars).

However, during TAP, the addition of a blocking mechanism dramatically increases the transcriptional noise (Figure 3F). To investigate the source of such high transcriptional noise during TAP, we simulated the timeseries of non-activated DNA and mRNA at the value of 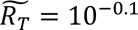 in TAP (Figure 3C, gray star). In the case of sole sequestration, non-activated DNA is rapidly restored to the activated state (Figure 3G, top left panel), and thus the timeseries of mRNA maintain their high average during TAP (Figure 3G, bottom left panel). In contrast, when blocking is added, this undesired repression during TAP persists for a long period (Figure 3G, top right panel), causing a dramatic reduction in the number of mRNAs (Figure 3G, bottom right panel). This significant reduction arises when the repressor, released from the activator due to noise, binds to the DNA-bound activator during TAP (unexpected repression; Figure 3H). Then, the repressor remains bound because there is no mechanism to release from the DNA-bound-activator until it will ultimately release itself (Figure 3H). This causes a dramatic reduction in the number of mRNAs even during TAP, leading to a small additional peak around zero in the mRNA distribution (Figure 3I). Due to the bimodality, the transcriptional noise surges during TAP (Figure 3F).

### A sequestration-, blocking-, and displacement-based switch is robust to noise

In the presence of blocking, the undesired repression during TAP stems from the unwanted binding of the repressor to the activator on DNA. This could be prevented by the addition of a displacement mechanism, which facilitates dissociation of this repressor-activator complex from the DNA. To investigate this, we incorporated displacement into our blocking and sequestration model. In this model, the repressor-activator complex (*R*_*A*_) can dissociate from the DNA with a dissociation constant of *K*_*d*_ (Figure 4A, gray box). We derived the transcriptional activity as the function of the molar ratio 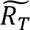 (Table S1; see Materials and Methods for details). The transcriptional activity can exhibit an ultrasensitive response to changes in 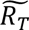 with the effective Hill coefficient of about 200 (Figure 4B, blue and red lines).

**Figure 4.**
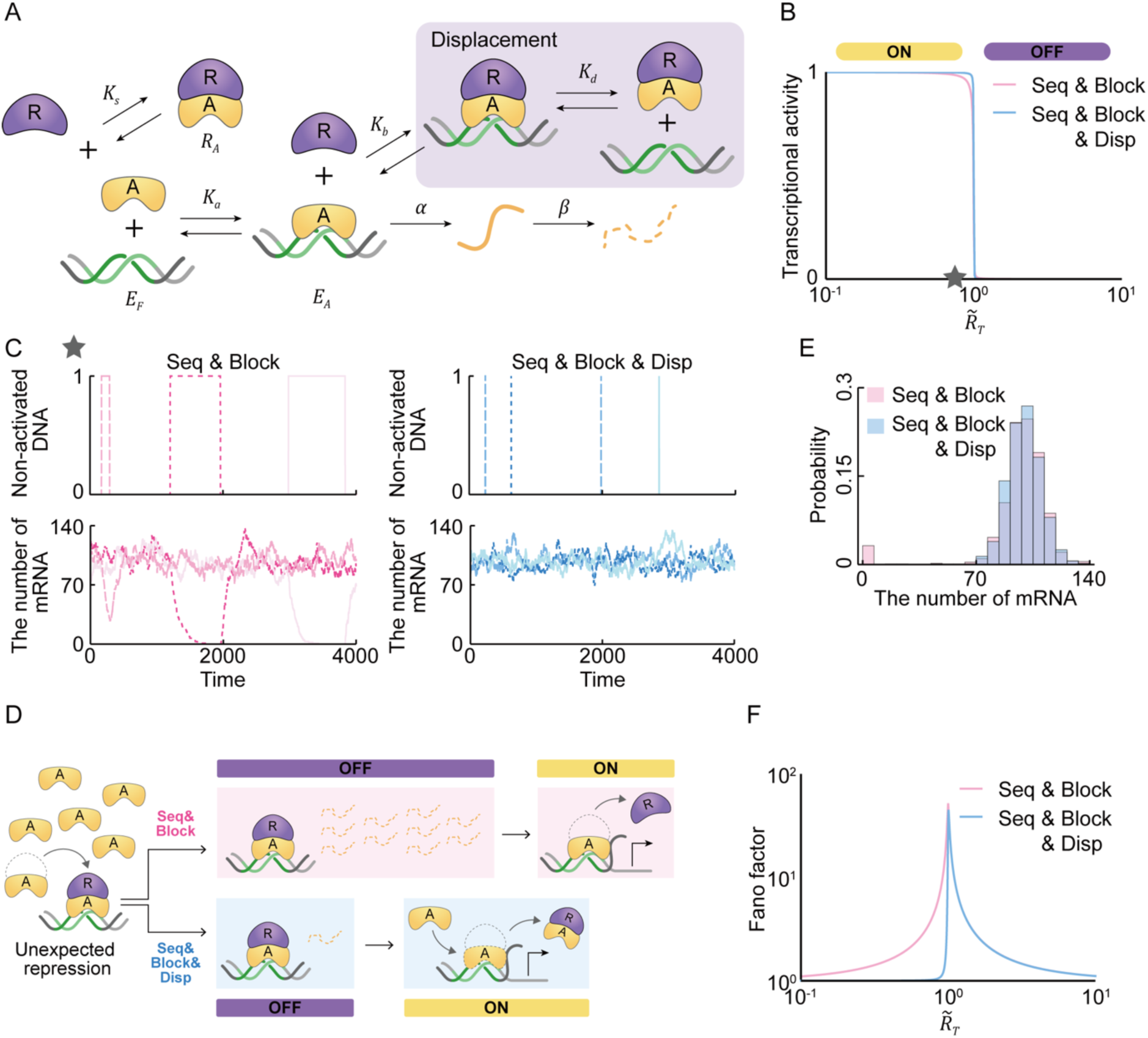
A transcriptional ultrasensitive switch based on the combination of sequestration, blocking, and displacement is robust to noise during both TAP and TRP. (**A**) Displacement is added to the blocking and sequestration model in Figure 3A. The repressor-activator complex (*R*_*A*_) dissociates from DNA (*E*_*d*_) with a dissociation constant of *K*_*d*_ to inhibit transcription (i.e., displacement; gray box). (**B**) The combination of sequestration, blocking and displacement (blue line) generates an ultrasensitive switch similar to sequestration and blocking (red line). (**C**) With the addition of displacement, the undesired repression of DNA during TAP is rapidly restored to the activated state. (**D**) When unwanted repression occurs during TAP, the repressor-activator complex dissociates from DNA and in the presence of displacement the free activator immediately promotes transcription (lower arrow), but not in its absence (upper arrow). (**E**) As a result, an additional peak around zero, observed with the combination of sequestration and blocking (red bars), is removed from the stationary distribution of mRNA (blue bars). (**F**) Thus, the addition of displacement leads to the reduction in the noise level during TAP.

Furthermore, the addition of displacement rapidly restores the undesired repression of DNA during TAP to the activated state (Figure 4C, top panel). This is evident in the simulated timeseries of mRNA, whose levels do not drop to zero (Figure 4C, bottom right panel) unlike with the combination of sequestration and blocking. That is, when a repressor binds to the DNA-bound-activator due to noise during TAP (unexpected repression; Figure 4), the repressor-activator complex is immediately released from DNA via displacement (Figure 4D, lower arrow). This does not happen without displacement (Figure 4D, upper arrow). Taken together, the inclusion of displacement eliminates the additional peak near zero seen in the sequestration and blocking model (Figure 4E), leading to a reduction in transcriptional noise during TAP (Figure 4F and Table S1).

## DISCUSSION

Traditionally, ultrasensitive transcriptional responses are generated by cooperative binding of transcriptional repressors to multiple DNA sites. This mechanism can trigger a sharp transition from transcriptional repression during TRP to activation during TAP as the number of repressors surpasses a threshold. However, the presence of noise can lead to prolonged periods of undesired repression during TAP and undesired activation during TRP (Figure 1). Thus, we shifted our focus from direct repression—where repressors directly bind to DNA, as seen in cooperative binding to multiple DNA sites—to the realm of indirect transcriptional repression: sequestration, blocking, and displacement. Sequestration alone can generate an ultrasensitive response that is attuned to changes in the molar ratio between repressors and activators. Specifically, when the number of repressors falls short of the number of activators (i.e., the molar ratio is less than one), an unsequestered activator binds to DNA, thereby promoting transcription. As the molar ratio surpasses one, a majority of activators become sequestered by repressors, promoting a sharp transition to transcriptional inhibition (Figure 2). Furthermore, the transcriptional repression (activation) during TRP (TAP) can be stably maintained by blocking (displacement) even in the presence of noise. Specifically, when an activator that is unexpectedly released from repressors binds to DNA and initiates transcription during TRP (Figure 2), the activator is immediately blocked by a repressor, resulting in the prompt restoration of transcriptional repression (Figure 3). When a repressor unexpectedly blocks an activator that is already bound to DNA during TAP, the repressor-activator complex is displaced, swiftly reinitiating transcriptional activation (Figure 4). Taken together, our exploration has unveiled an ultrasensitive transcriptional switch endowed with resilience against noise during both TAP and TRP. This breakthrough holds the potential to enhance the stability and reliability of transcriptional regulation within intricate biological systems. Furthermore, it would be interesting future work to explore extensions to a bistable switch (29).

Interestingly, although two models with or without a particular repression mechanism may appear similar in deterministic simulations, they can behave quite differently in stochastic simulations. Specifically, in the deterministic ODE, where the transcription of mRNA is proportional to the transcriptional activity, two models exhibiting similar transcriptional activity yield indistinguishable behaviors of mRNA (18). On the other hand, in the stochastic models, the distributions of mRNA can be completely different despite resembling transcriptional activities. For example, all transcriptional activity of each model shows similar responses with respect to changes in the molar ratio (Figures 3B and 4B), whereas their Fano factors are completely different (Figures 3F and 4F). Therefore, a comprehensive understanding of the precise repression mechanism is essential to ensure unbiased modeling results when considering the presence of noise.

Biological oscillators and biological switches require ultrasensitivity to generate strong rhythms (28,30) and bistable responses (2,31), respectively. In addition to ultrasensitivity, these systems need to exhibit robustness against intrinsic noise to ensure accurate timing and precise decision-making. Indeed, such biological systems that require both ultrasensitivity and precision employ multiple indirect repression mechanisms. For instance, the NF-κB oscillator is known to utilize both sequestration and displacement to inhibit NF-κB by IκK (17,19) (Figure 5A). The p53-MDM2 oscillator (32,33) (Figure 5B), and circadian clocks in mammals (Figure 5C) (21-23) and Drosophila (20,34) employ a combination of sequestration, blocking, and displacement. Notably, in the mammalian circadian clock, disruption of displacement by the CK18^Δ2/Δ2^ mutant leads to an increase in the variance of peak-to-peak periods of circadian rhythms (Figure 5D). This result indicates that the combination of sequestration, blocking, and displacement is required for precise on and off of transcriptional negative loop in the mammalian circadian clock. Our work highlights the significance of multiple repression mechanisms in achieving robust and precise regulatory outcomes in these cellular systems.

**Figure 5.**
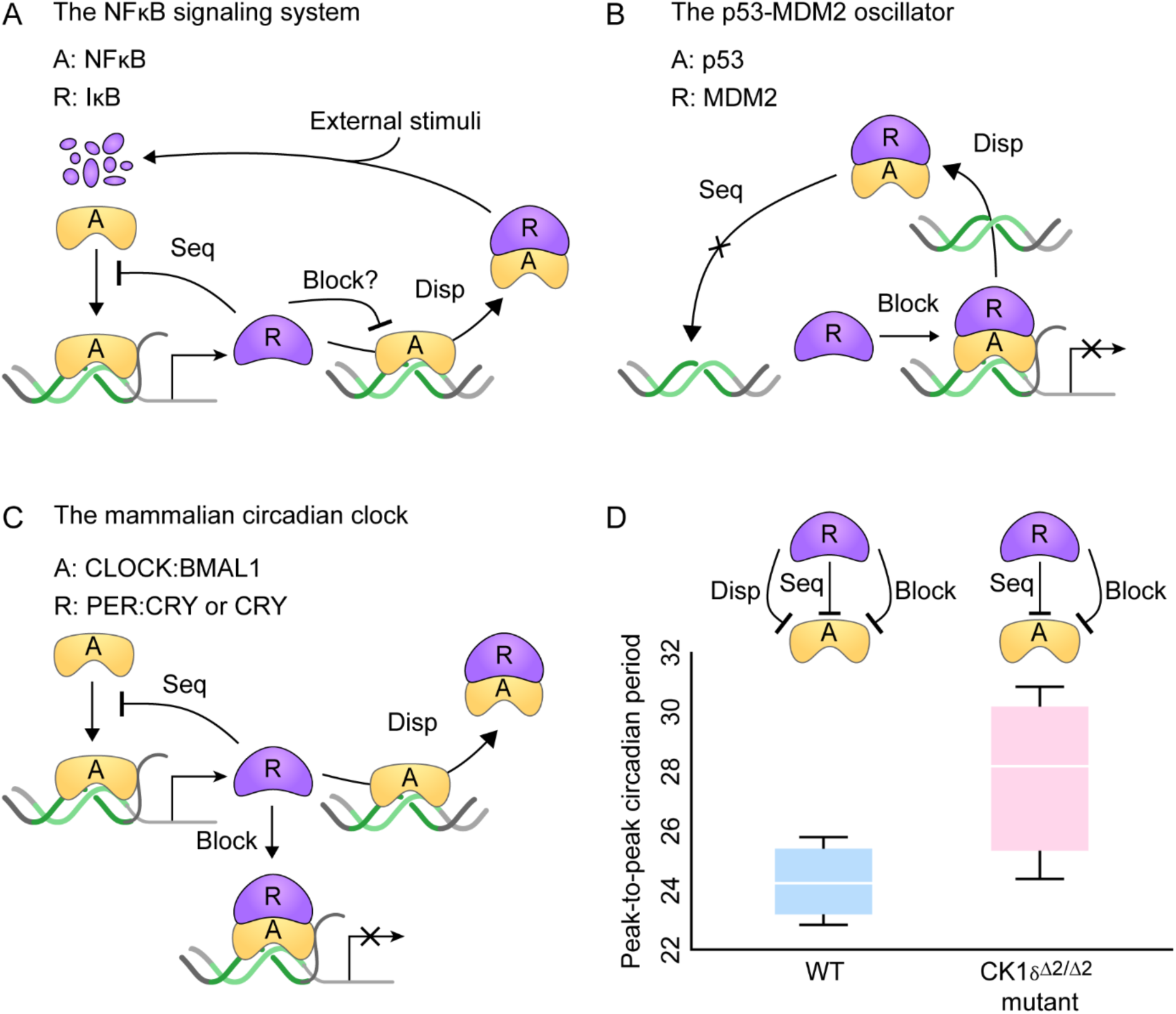
Various biological systems, requiring a precise ultrasensitive switch, utilize the combination of sequestration, blocking, and displacement. (**A**) In the NF-κB oscillator, IκB can repress stimulation-induced transcription by displacing the transcriptional activator NF-κB from DNA, as well as by sequestering it in the cytoplasm. It has not been investigated whether IκB can block the transcription by binding to DNA-bound-NF-κB. (**B**) In the p53-MDM2 oscillator, MDM2 binds with p53 to displace from DNA, and then sequesters it not to bind with DNA. Furthermore, MDM2 can block the transcriptional activity of p53 with a corepressor. (**C**) In the mammalian circadian clock, the complex of PER and CRY inhibits their own transcription by sequestering and displacing the transcriptional activator CLOCK:BMAL1, while CRY inhibits the transcription by binding solely to the CLOCK:BMAL1:DNA complex. (**D**) When the displacement is disrupted by the CK181′2/1′2 mutant in the mammalian circadian clock, the variance of peak-to-peak periods of circadian rhythms increases (WT: 24.0 ± 1.6 h, CK18^Δ2/Δ2^: 27.9 ± 3.6 h). The peak-to-peak period is measured using data retrieved from Etchegaray *et al*.

Currently existing synthetic switches have been designed based on direct DNA-binding mechanisms such as cooperative binding on multiple binding sites (35), molecular titration via decoy binding sites (6), and multistage transcriptional cascade (36), or based on a single indirect repression mechanism, i.e., sequestration (25,26). When these mechanisms generate ultrasensitivity, they become sensitive to noise. For instance, as super enhancers activated via the cooperative binding of the transcriptional activator generate a more sensitive transcriptional response, the variance of mRNA distribution becomes larger (37). Furthermore, as the binding affinity between the transcription factor and the decoy sites becomes stronger, the switch becomes more sensitive, but the noise level in transcription also increases (9). Similarly, as the number of stages of cascades increases, the sensitivity of the switch increases, but the noise level in transcription also increases (36). Furthermore, previous studies have also pointed out that a single sequestration-based switch can amplify noise (26), consistent with our results (Figure 2G). On the other hand, when additional indirect transcriptional repression mechanisms are added, while the sensitivity of transcriptional response increases, the sensitivity to noise decreases. Thus, the combination of the diverse indirect transcriptional repression mechanisms proposed in this study provides a design for synthetic ultrasensitive switches that are also robust to noise.

## DATA AVAILABILITY

The underlying code for this study is not publicly available but may be made available to qualified researchers on reasonable request.

## SUPPLEMENTARY DATA

Supplementary Data are available at NAR online.

## AUTHOR CONTRIBUTIONS

Eui Min Jeong: Conceptualization, Formal analysis, Methodology, Visualization, Writing— original draft. Jae Kyoung Kim: Conceptualization, Formal analysis, Supervision, Visualization, Writing—original draft.

## ACKNOWLEDGEMENTS

The authors thank Life Science Editors for editing support.

## REFERENCES

1. Ferrell, J.E., Jr. and Ha, S.H. (2014) Ultrasensitivity part I: Michaelian responses and zero-order ultrasensitivity. Trends Biochem Sci, 39, 496–503.

2. Ferrell, J.E., Jr. and Ha, S.H. (2014) Ultrasensitivity part III: cascades, bistable switches, and oscillators. Trends Biochem Sci, 39, 612–618.

3. Zhang, Q., Bhattacharya, S. and Andersen, M.E. (2013) Ultrasensitive response motifs: basic amplifiers in molecular signalling networks. Open Biol, 3, 130031.

4. Mirny, L.A. (2010) Nucleosome-mediated cooperativity between transcription factors. Proc Natl Acad Sci U S A, 107, 22534–22539.

5. Lu, M.S., Mauser, J.F. and Prehoda, K.E. (2012) Ultrasensitive synthetic protein regulatory networks using mixed decoys. ACS Synth Biol, 1, 65–72.

6. Lee, T.H. and Maheshri, N. (2012) A regulatory role for repeated decoy transcription factor binding sites in target gene expression. Mol Syst Biol, 8, 576.

7. Riccione, K.A., Smith, R.P., Lee, A.J. and You, L. (2012) A synthetic biology approach to understanding cellular information processing. ACS Synth Biol, 1, 389–402.

8. Sanchez, A., Garcia, H.G., Jones, D., Phillips, R. and Kondev, J. (2011) Effect of promoter architecture on the cell-to-cell variability in gene expression. PLoS Comput Biol, 7, e1001100.

9. Dey, S., Soltani, M. and Singh, A. (2020) Enhancement of gene expression noise from transcription factor binding to genomic decoy sites. Sci Rep, 10, 9126.

10. Shibata, T. and Fujimoto, K. (2005) Noisy signal amplification in ultrasensitive signal transduction. Proc Natl Acad Sci U S A, 102, 331–336.

11. Gaston, K. and Jayaraman, P.S. (2003) Transcriptional repression in eukaryotes: repressors and repression mechanisms. Cell Mol Life Sci, 60, 721–741.

12. Renkawitz, R. (1990) Transcriptional repression in eukaryotes. Trends Genet, 6, 192–197.

13. Benezra, R., Davis, R.L., Lockshon, D., Turner, D.L. and Weintraub, H. (1990) The protein Id: a negative regulator of helix-loop-helix DNA binding proteins. Cell, 61, 49–59.

14. *Van Doren*, M., *Ellis*, H.M. and *Posakony*, J.W. (1991) The Drosophila extramacrochaetae protein antagonizes sequence-specific DNA binding by daughterless/achaete-scute protein complexes. Development, 113, 245–255.

15. Chadsey, M.S., Karlinsey, J.E. and Hughes, K.T. (1998) The flagellar anti-sigma factor FlgM actively dissociates Salmonella typhimurium sigma28 RNA polymerase holoenzyme. Genes Dev, 12, 3123–3136.

16. Jayaraman, P.S., Hirst, K. and Goding, C.R. (1994) The activation domain of a basic helix-loop-helix protein is masked by repressor interaction with domains distinct from that required for transcription regulation. EMBO J, 13, 2192–2199.

17. Potoyan, D.A., Zheng, W., Komives, E.A. and Wolynes, P.G. (2016) Molecular stripping in the NF-kappaB/IkappaB/DNA genetic regulatory network. Proc Natl Acad Sci U S A, 113, 110–115.

18. Jeong, E.M., Song, Y.M. and Kim, J.K. (2022) Combined multiple transcriptional repression mechanisms generate ultrasensitivity and oscillations. Interface Focus, 12, 20210084.

19. Bergqvist, S., Alverdi, V., Mengel, B., Hoffmann, A., Ghosh, G. and Komives, E.A. (2009) Kinetic enhancement of NF-kappaBxDNA dissociation by IkappaBalpha. Proc Natl Acad Sci U S A, 106, 19328–19333.

20. Menet, J.S., Abruzzi, K.C., Desrochers, J., Rodriguez, J. and Rosbash, M. (2010) Dynamic PER repression mechanisms in the Drosophila circadian clock: from on-DNA to off-DNA. Genes Dev, 24, 358–367.

21. Cao, X., Yang, Y., Selby, C.P., Liu, Z. and Sancar, A. (2021) Molecular mechanism of the repressive phase of the mammalian circadian clock. Proc Natl Acad Sci U S A, 118.

22. Chiou, Y.Y., Yang, Y., Rashid, N., Ye, R., Selby, C.P. and Sancar, A. (2016) Mammalian Period represses and de-represses transcription by displacing CLOCK-BMAL1 from promoters in a Cryptochrome-dependent manner. Proc Natl Acad Sci U S A, 113, E6072–E6079.

23. Ye, R., Selby, C.P., Chiou, Y.Y., Ozkan-Dagliyan, I., Gaddameedhi, S. and Sancar, A. (2014) Dual modes of CLOCK:BMAL1 inhibition mediated by Cryptochrome and Period proteins in the mammalian circadian clock. Genes Dev, 28, 1989–1998.

24. Ferrell, J.E., Jr. and Ha, S.H. (2014) Ultrasensitivity part II: multisite phosphorylation, stoichiometric inhibitors, and positive feedback. Trends Biochem Sci, 39, 556–569.

25. Buchler, N.E. and Cross, F.R. (2009) Protein sequestration generates a flexible ultrasensitive response in a genetic network. Mol Syst Biol, 5, 272.

26. Buchler, N.E. and Louis, M. (2008) Molecular titration and ultrasensitivity in regulatory networks. J Mol Biol, 384, 1106–1119.

27. Kim, J.K. and Forger, D.B. (2012) A mechanism for robust circadian timekeeping via stoichiometric balance. Mol Syst Biol, 8, 630.

28. Kim, J.K. (2016) Protein sequestration versus Hill-type repression in circadian clock models. IET Syst Biol, 10, 125–135.

29. Nordick, B., Yu, P.Y., Liao, G. and Hong, T. (2022) Nonmodular oscillator and switch based on RNA decay drive regeneration of multimodal gene expression. Nucleic Acids Res, 50, 3693–3708.

30. Novak, B. and Tyson, J.J. (2008) Design principles of biochemical oscillators. Nat Rev Mol Cell Biol, 9, 981–991.

31. Rombouts, J. and Gelens, L. (2021) Dynamic bistable switches enhance robustness and accuracy of cell cycle transitions. PLoS Comput Biol, 17, e1008231.

32. Cross, B., Chen, L., Cheng, Q., Li, B., Yuan, Z.M. and Chen, J. (2011) Inhibition of p53 DNA binding function by the MDM2 protein acidic domain. J Biol Chem, 286, 16018–16029.

33. Chen, J. (2016) The Cell-Cycle Arrest and Apoptotic Functions of p53 in Tumor Initiation and Progression. Cold Spring Harb Perspect Med, 6, a026104.

34. Jeong, E.M., Kwon, M., Cho, E., Lee, S.H., Kim, H., Kim, E.Y. and Kim, J.K. (2022) Systematic modeling-driven experiments identify distinct molecular clockworks underlying hierarchically organized pacemaker neurons. Proc Natl Acad Sci U S A, 119.

35. Dueber, J.E., Mirsky, E.A. and Lim, W.A. (2007) Engineering synthetic signaling proteins with ultrasensitive input/output control. Nat Biotechnol, 25, 660–662.

36. Hooshangi, S., Thiberge, S. and Weiss, R. (2005) Ultrasensitivity and noise propagation in a synthetic transcriptional cascade. Proc Natl Acad Sci U S A, 102, 3581–3586.

37. Michida, H., Imoto, H., Shinohara, H., Yumoto, N., Seki, M., Umeda, M., Hayashi, T., Nikaido, I., Kasukawa, T., Suzuki, Y. et al. (2020) The Number of Transcription Factors at an Enhancer Determines Switch-like Gene Expression. Cell Rep, 31, 107724.

